# Reagent-free Raman and quantitative phase imaging offer a unique morpho-molecular platform for recognition of malignancy and stages of B-cell acute lymphoblastic leukemia

**DOI:** 10.1101/2021.01.25.428006

**Authors:** Santosh Kumar Paidi, Piyush Raj, Rosalie Bordett, Chi Zhang, Sukrut Hemant Karandikar, Rishikesh Pandey, Ishan Barman

## Abstract

Acute lymphoblastic leukemia (ALL) is one of the most common malignancies which account for nearly one-third of all pediatric cancers. The current diagnostic assays are time-consuming, labor-intensive, and require expensive reagents. Here, we report a label-free approach featuring diffraction phase imaging and Raman microscopy that can retrieve both morphological and molecular attributes for label-free optical phenotyping of individual B cells. By investigating leukemia cell lines of early and late stages along with the healthy B cells, we show that phase image can capture subtle morphological differences among the healthy, early, and late stages of leukemic cells. By exploiting its biomolecular specificity, we demonstrate that Raman microscopy is capable of accurately identifying not only different stages of leukemia cells, but also individual cell lines at each stage. Overall, our study provides a rationale for employing this hybrid modality to screen leukemia cells using the widefield QPI and using Raman microscopy for accurate differentiation of early and late-stage phenotypes. This contrast-free and rapid diagnostic tool exhibits great promise for clinical diagnosis and staging of leukemia in the near future.

## Introduction

Acute lymphoblastic leukemia (ALL) is one of the most common cancers in pediatric population due to malignant clonal proliferation of lymphoid progenitor cells. B-cell ALL (B-ALL) is the leading cause of cancer-related deaths in children and adolescents [1]. The clinical diagnosis of B-ALL currently relies on the use of multi-parametric flow cytometry for the identification and classification of B-ALL. However, flow cytometry-based assays require expensive high-quality antibodies and extensive sample preparation; additionally, non-specific signals due to antibody cross-reactivity and photobleaching pose challenges. Several other methods such as those based on mass cytometry [2], DNA methylation [3], and gene expression [4] are also characterized by some of these limitations. Therefore, there is an unmet need for analytical tools that can provide a facile route for quantitative and early diagnosis and staging of B-ALL.

In this milieu, techniques based on label-free optical imaging and molecular spectroscopy are emerging as attractive alternatives for cancer cell phenotyping based on their morphological and biomolecular information content [5–11]. Specifically, quantitative phase imaging (QPI) has been used to investigate morphology of live cells and identify disease-specific cellular phenotypes [12–22]. On the other hand, Raman spectroscopy has been employed to measure the biomolecular composition with single cell resolution for cell differentiation and disease identification [23–32]. Since both the molecular and morphological changes are associated with malignant transformations, we have sought to develop a morphomolecular microscopy (3M) platform, which combines the complementary morphological and biochemical information content into diagnostic frameworks [33, 34]. There are two main motivations to combine Raman spectroscopy and QPI. First, the increased information content of this hybrid modality confers diagnostic advantage by accounting for alterations in both the cellular attributes due to disease onset and progression. Second, the overall sampling time of a Raman measurement can be reduced by imaging only the suspicious cell populations by leveraging high-throughput characteristics of QPI for coarse cellular analysis or screening. In this study, we employed our custom-built 3M system that leverages the widefield imaging capability of QPI and exquisite molecular specificity of Raman spectroscopy to derive complementary data from a panel of B-ALL cells at different stages (**Fig. 1**) [33]. Using visually aided morpho-phenotyping recognition (VAMPIRE) tool, we reveal that the progression of B-ALL is characterized by robust shape mode changes captured by QPI [35, 36]. We also show that the Raman spectroscopic data provided adequate clustering capability based on the cell type in principal component analysis-derived component space. Furthermore, the supervised classification routine based on random forest analysis enabled accurate classification of the B-ALL cells based on both the stage of disease progression and the cell line identity. Such an approach, fusing the biomolecular analysis afforded by Raman spectroscopy with morphological assessment capability of rapid widefield QPI, could be exploited to accelerate the phenotype recognition of single cells by sampling/screening a subset of cells based on their biophysical properties for downstream biomolecular characterization.

**Figure 1.**
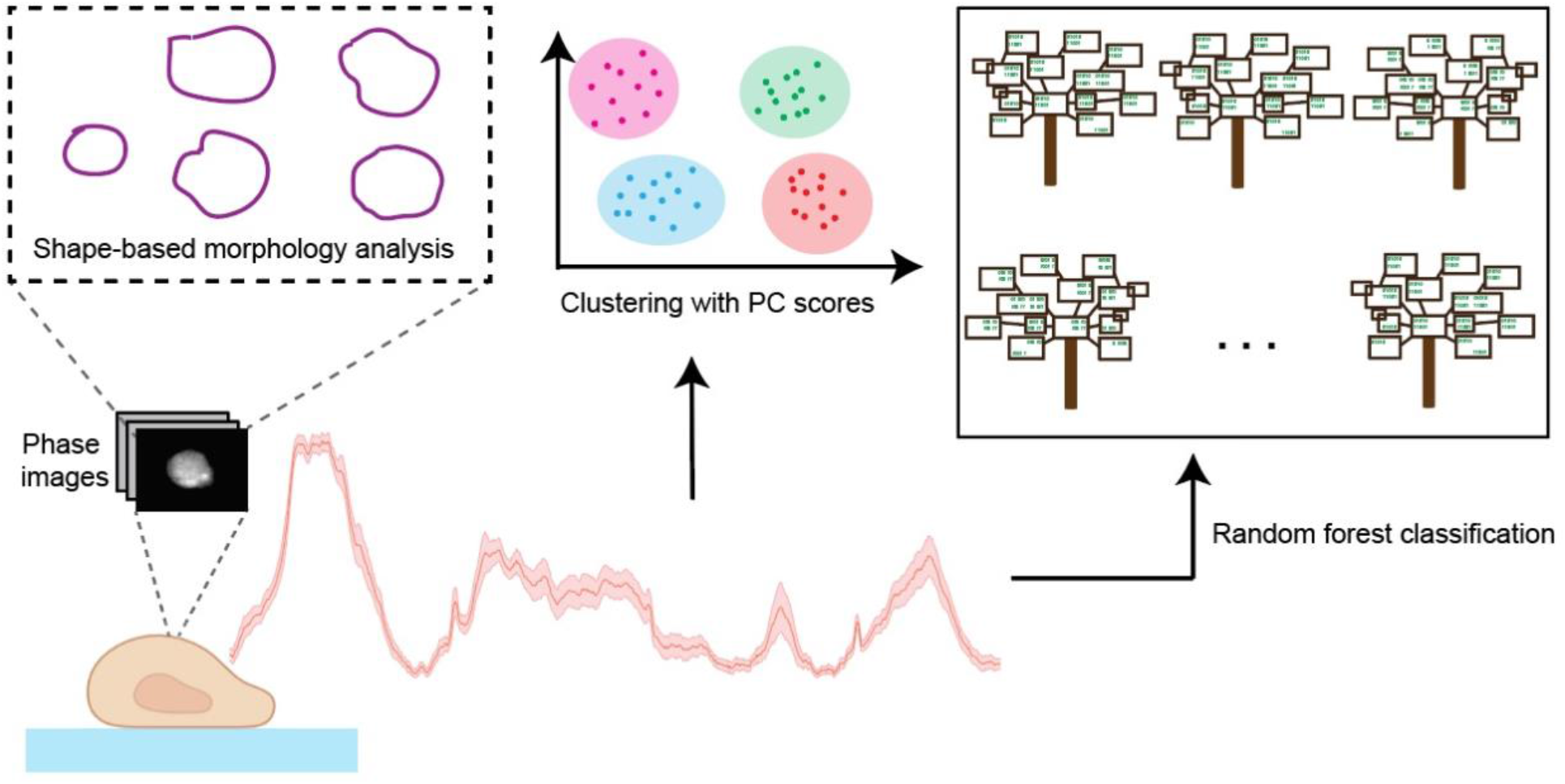
Study overview. Morphomolecular assessment of B-ALL using Raman spectroscopy and quantitative phase imaging.

## Results and discussion

We used leukemic B cells from early (REH, RS4;11) and late (BALL-1, MN60) stages of malignancy along with healthy B-cells (Control) for morpho-molecular characterization using QPI and Raman spectroscopy. **Figure 2A** illustrates the expression of various known surface markers to assess the purity of the cell lines. We also determined the percentage enrichment of healthy B cells from the isolated B cells (**Fig. 2B**). QPI images of cells from REH (n = 389), RS4;11 (n = 406), BALL-1 (n = 394), MN60 (n = 415), and Control (n = 690) classes were acquired (**Fig. 3A**). To characterize morphological changes associated with B-ALL progression, we employed the recently developed VAMPIRE tool to analyze QPI images [35, 36]. The VAMPIRE tool identifies differences in morphological phenotype in terms of the distribution of the cell population in each class into few representative shape modes determined by dimensionality reduction of morphological parameters [35, 36]. By subjecting the entire QPI dataset comprising of 2294 images to the VAMPIRE analysis, we determined five unique shape modes that capture variance in cellular morphology across the dataset (**Fig. 3B**). The distribution of cells into these shape modes was determined by K-means clustering to identify the representative shape mode for each cell in the dataset. We observed distinct patterns of the cell shape mode distributions for the cells that is indicative of the differences in the stages of B-ALL progression. While the distributions for the Control group is skewed towards shape modes 1 and 2 and the distributions for the late-stage cells are skewed towards shape modes 4 and 5, the distributions for early-stage cell groups are centered around the middle shape modes marking a transition between the healthy and late-stage cells (**Fig. 3B**). While the shape modes constitute abstract transformations of conventional morphological descriptors, analysis of these shape modes clearly reveals that there exist characteristics of B-ALL progression that are directly related to the morphological attributes of these cell populations. Furthermore, the robustness of such abstract markers based on cell morphology is confirmed by the similarity of the distributions for both the cell lines within each stage-specific group of B-ALL progression. On the other hand, this similarity makes the discrimination between cell types at similar stages based solely on morphological parameters challenging.

**Figure 2.**
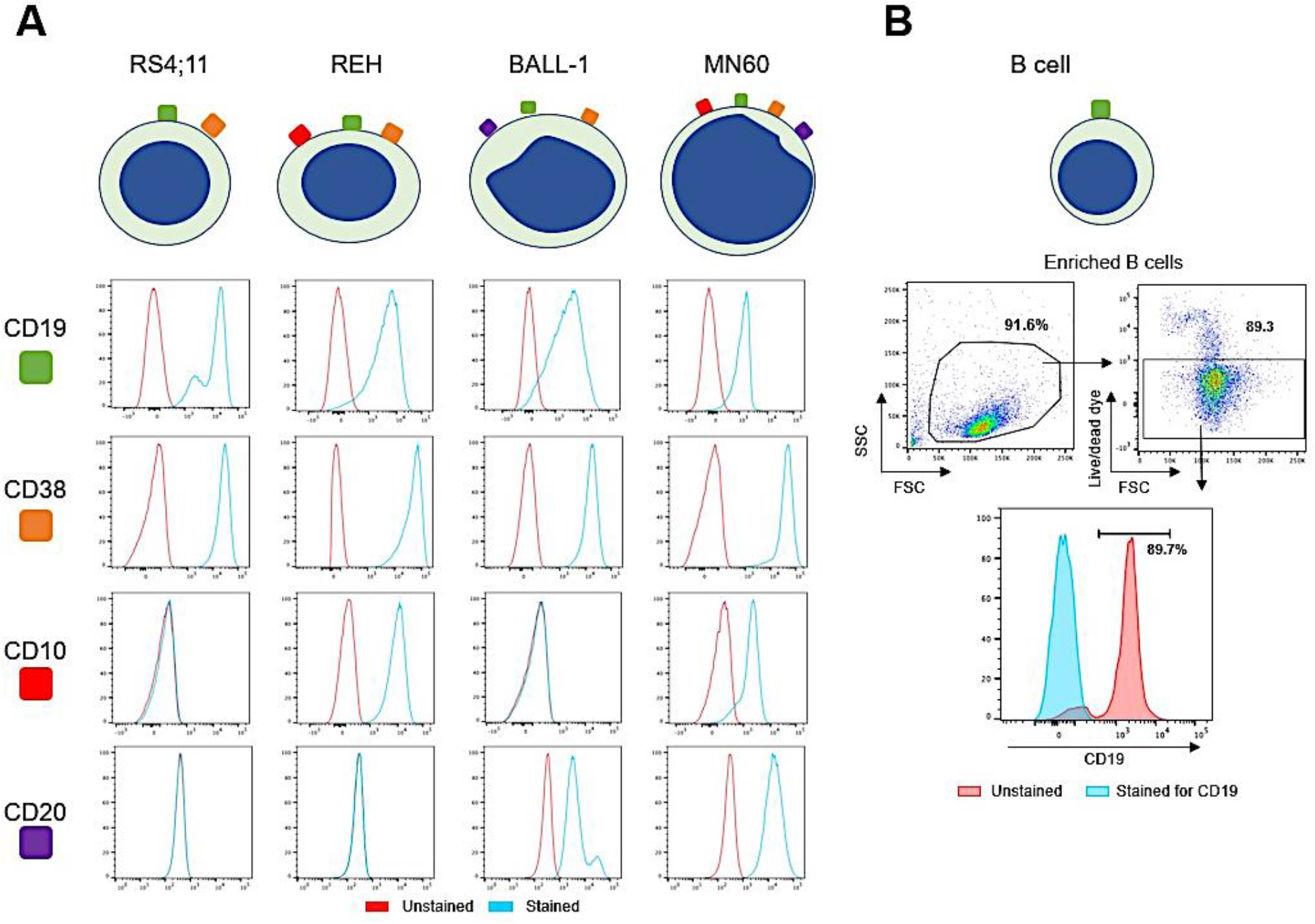
Flow cytometry characterization of B-ALL cell lines and control healthy B-cells. **(A)** The expression profiles of characteristic surface markers of B-cells are shown for the cell lines used in the study. **(B)** The percentage enrichment of healthy B-cells from the isolated B-cells as assessed using flow cytometry.

**Figure 3.**
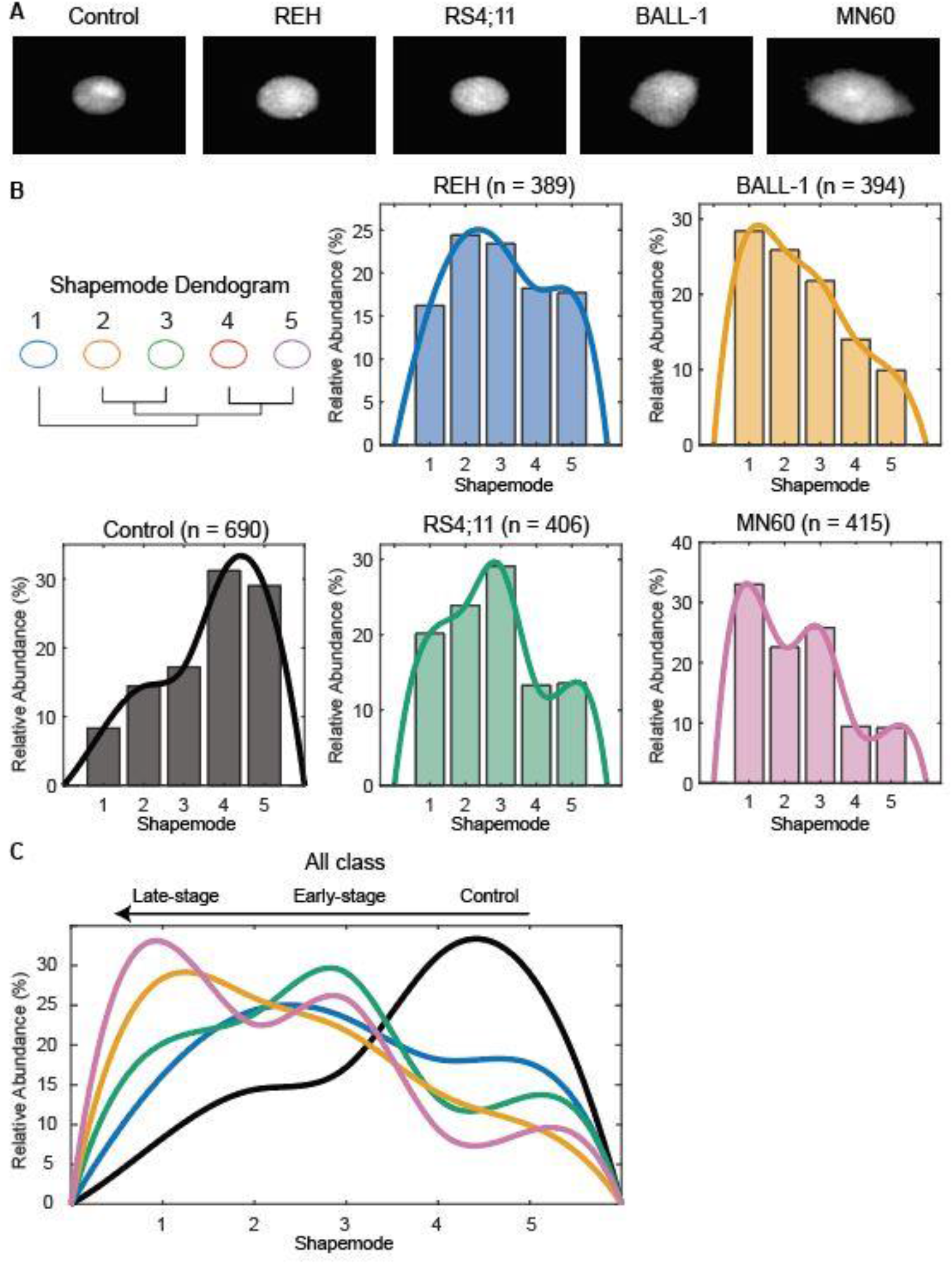
Quantitative phase imaging of B-ALL. **(A)** Representative phase images of the Control, early-stage (REH and RS4;11), and late-stage (BALL-1 and MN60) B-ALL cells are shown. **(B)** The histograms show the relative abundance profiles of the five shape modes identified using VAMPIRE analysis in each cell group. The overlapping curves fitted to the histogram show the skew of the distribution of the shape modes in each cell group. **(C)** The comparison of the fitted curves shows the progression of the abundant shape mode from right to left in response to malignant progression of B-ALL.

To tackle this limitation of morphological analysis, we employed Raman spectroscopy to complement the morphological information captured by QPI with rich molecular information indicative of chemical makeup of the B-ALL cells. We used our custom-built Raman microscope to collect a total of 959 Raman spectra from cells in Control, REH, RS4;11, BALL-1, and MN60 classes. The mean and standard deviation of the spectra in each class are shown in **Fig. 4A**. The spectra in the fingerprint region exhibit peaks representative of biological components of cells, particularly at 790 cm^−1^ (O-P-O stretching of DNA), 1008 cm^−1^ (C–C stretching vibration of phenylalanine), 1125 cm^−1^ (C-C stretching mode of lipids and C-N stretch of proteins), 1452 cm^−1^ (CH_2_ bending modes in lipids and proteins) and 1654 cm^−1^ (amide I of proteins and C=C stretching in lipids) [37]. We performed principal component analysis of the dataset to perform dimensionality reduction and obtain a set of components ordered according to the variance in the spectral dataset they capture (**Fig. S1, Supporting Information**). Each spectrum in the dataset is described in terms of these component loading vectors and the corresponding scores. We used the scores of the PC loadings to investigate if specific clustering patterns exist in the spectral dataset that correlate with the stage of the disease progression. Using Radviz radial visualization tool in Orange datamining software, we plotted the scores of the component loadings that maximize the separation of the clusters based on the cell type (**Fig. 4B**) [38]. We observed appreciable clustering of spectra from each cell type and localization of the clusters belonging to cells in the same stage of progression on the Radviz score map. The larger overlap of the early-stage REH and RS4;11 clusters in comparison to the late-stage BALL-1 and MN60 clusters hints at the divergence in molecular composition between different leukemia cells stemming from further disease progression. However, the reasonable separation of both early-stage and late-stage cell clusters from the Control cluster indicate that Raman spectroscopy can capture subtle molecular changes associated with leukemia – even during the early stages of transformation.

**Figure 4.**
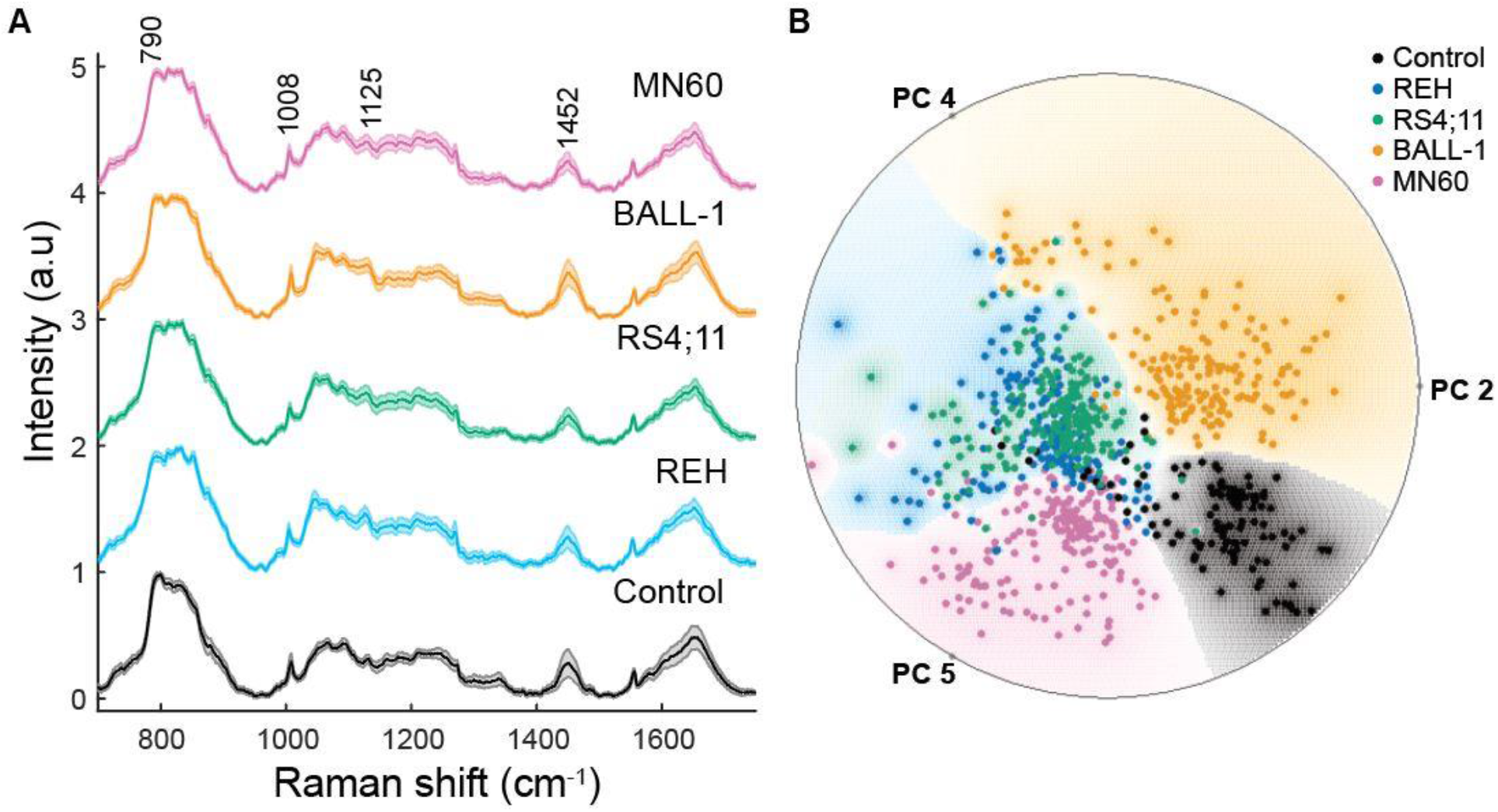
Raman spectroscopy of B-ALL. **(A)** Shaded error plots show the mean and standard deviation of the Raman spectra obtained from cells in each group. The key spectral features of the spectra are highlighted. **(B)** The radial visualization plot shows the clustering of the Raman spectra obtained from the Control and B-ALL cells in the principal component space.

Motivated by these promising results, we sought to employ supervised classification techniques for quantitative assessment of Raman spectroscopy for identification of the leukemia cells. Therefore, we employed random forests – a method based on ensemble of decision trees – for deriving classifiers for the present application. We performed multiclass random forest analysis by using the identity of the five cell lines as the class labels for the spectra. Specifically, we trained five-class random forest classifiers composed of 100 decision trees and computed the out-of-bag classification error, which is a measure of performance based on testing the classifiers using samples that have been left out of training. To further improve the robustness of the classification results, we performed 200 iterations of random forest classification by including random subsets of spectra from each class to ensure equal representation of all the classes in the model training. We observed that the out-of-bag classification error decreased below 5% through inclusion of ~65 trees for the five-class classification (**Fig. 5A**). We also found that the high accuracy of classification is consistent for all the five cell lines (**Fig. 5B**). While the high classification accuracy levels underscore that the acquired Raman spectra encode subtle, yet reproducible, differences in the biomolecular composition of various malignant leukemia cell types, it is also important to verify that the developed classifiers can be used to identify the stage of disease progression. Therefore, we performed the same random forest analysis by using the stage of leukemia progression as an outcome to train a three-class model for classification of control, early-stage, and late-stage cells. The out-of-bag error rate for the three-class classification based on the stage of the cells in leukemia progression also showed similar decline to below 5% by including only 65 trees (**Fig. 5C**). The accuracy for recognition of individual classes also show that supervised classification based on Raman spectra is able to accurately distinguish healthy B-cells from early-stage and late-stage leukemia cells (**Fig. 5D**).

**Figure 5.**
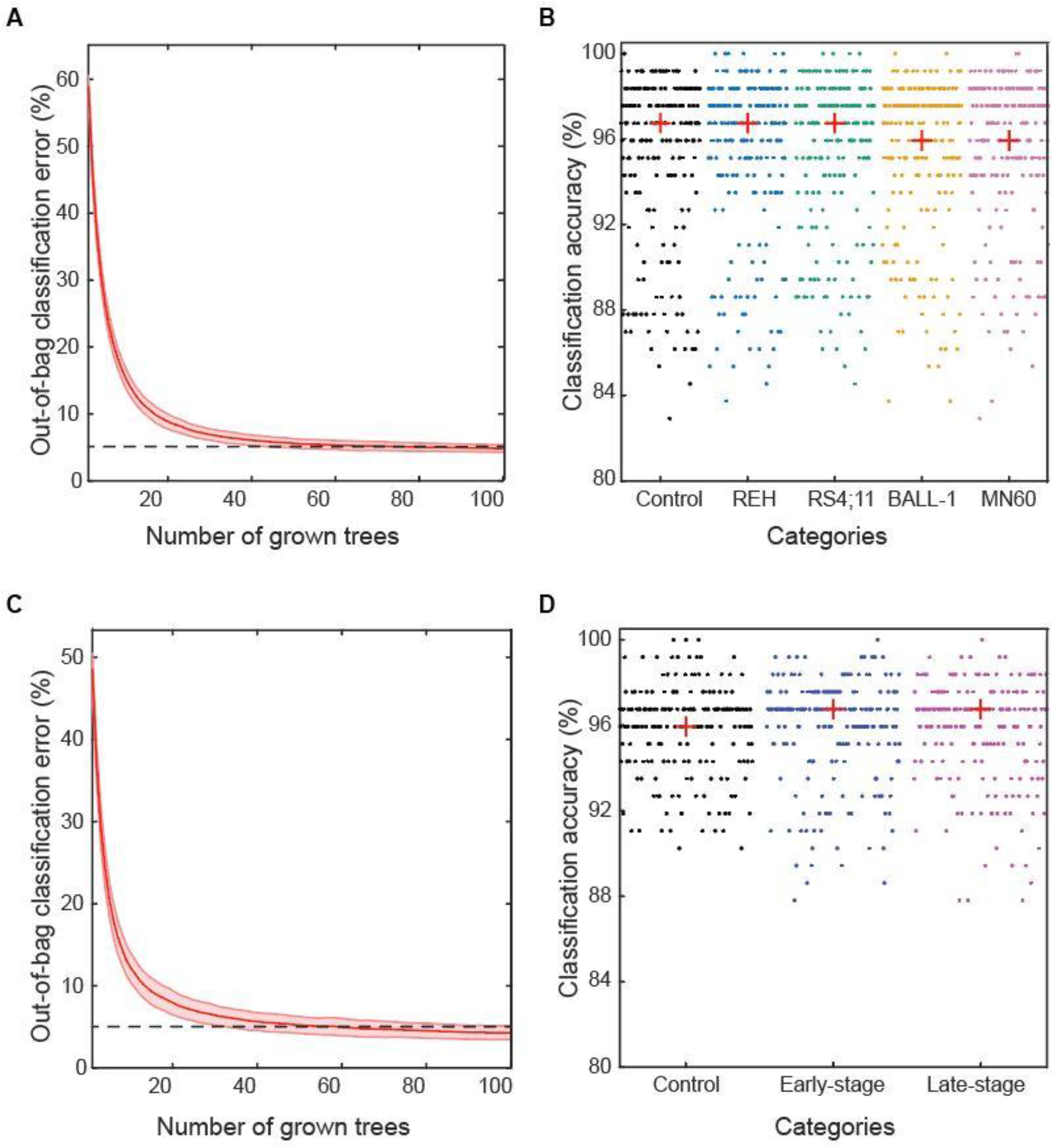
Random forest classification of Raman spectra. **(A)** The out-of-bag classification error plot, which illustrates the evolution of classification error as a function of number of decision trees included in the forest, shows that the error for random forest classifiers trained on the Raman spectra from the five cell lines falls below 5% with the inclusion of less than 100 trees in the forest. The shadow represents standard deviation of error across 200 iterations. **(B)** The corresponding classification accuracy for each B-ALL cell line and healthy B-cells are plotted for all the 200 iterations. **(C)** The out-of-bag classification error plot shows the evolution of error for the random forest classifiers trained on the stage of the cells in B-ALL progression. **(D)** The corresponding accuracy for control, early-stage, and late-stage cells are shown for 200 iterations.

While these results showcase the promise of this versatile and widely accessible morpho-molecular platform, it is worth noting the limitation of the present proof-of-principle study. The QPI and Raman measurements have not been performed on the same set of cells thereby limiting our ability to infer the direct benefit of fusing these two datasets. Nevertheless, acquisitions of a significantly large number of images and spectra accounts for the cellular heterogeneity to a large extent through adequate sampling of the populations. Furthermore, the recent improvement in our system now enables sequential Raman and QPI measurements from the same cells allowing us to directly address this drawback in our future studies. Overall, our results show that while morphological characterization of the leukemia cells provides clear demarcation between the cells at various stages of disease progression, the rich molecular information captured by Raman spectroscopy can provide a route for additional demarcation between different cell groups at similar stages.

## Conclusions

Our proposed approach makes it possible to distinguish between healthy, early-stage and late-stage cells B cells and show the promise of this morpho-molecular platform for fully automated, label-free and rapid diagnosis of B-ALL. Importantly, a recent study from our laboratory indicates that QPI can be employed to discriminate B-cells from other leucocytes [39]. This would obviate the need of any additional separation method to select B-cells either from the whole blood or from the leucocyte mixtures. Consequently, leucocyte mixture, which can be very easily separated from the peripheral blood using centrifugation, can be directly used for the proposed hybrid assay. This fully label-free strategy provides a unique platform in which the malignant B-cells can be optically interrogated while also preserving them for subsequent cytogenetic and immunophenotypic analysis.

## Material and methods

### Healthy control B cells

The fresh blood samples from three anonymous healthy adult donors were purchased from StemCell Technologies (Vancouver, Canada) and all the experiments were conducted within 24 hours of blood donation. The purchased blood samples contained ethylenediaminetetraacetic acid (EDTA) as anti-coagulant. The B cells were isolated from fresh blood samples using negative selection kits from Stemcell Technologies (EasySep Direct Human B Cell Isolation Kit). This separation kits uses immunomagnetic negative selection for isolating B cells from the whole blood sample. The isolation was carried out following the manufacturer’s instructions and B cells were isolated in phosphate-buffered saline (PBS) free from Ca++ and Mg++ (Gibco, Thermo Fisher Scientific). The B cells suspended in the PBS from the final incubation were centrifuged at 400g for 5 minutes. The cell pellet was resuspended in PBS before the cells were imaged.

### Cell lines

We purchased the early-stage leukemia cell lines REH and RS4;11 from American Type Culture Collection (ATCC, USA) and procured the late-stage leukemia cell lines BALL-1 and MN60, from RIKEN BioResource Research Center (Japan) and DSMZ (Germany), respectively. All the four cell lines were cultured according to previously described protocols before imaging measurements [40].

### Flow cytometry

Both primary cells and cell lines were tested using multiparametric flow cytometry for their purity. Anti-CD10-PerCP/Cy5.5 (Cat no. 312215), anti-CD19-APC (Cat no.392503), anti-CD20-FITC (Cat no. 302303) and anti-CD38-PE (Cat no 356603) were purchased from Biolegend. Fluorescent viability dye, eFlour780, was purchased from ThermoFisher Scientific (Cat no. 65-0865-14). Cells were stained with individual antibodies (quantity as recommended by the manufacturer) in cold PBS containing 0.5% fetal calf serum and the fluorescent viability dye (1:1000 dilution) for 20 minutes. Cells were maintained at a density of one million cells per 100 ml of buffer. Cells were then washed three times with 200 μl of the buffer and resuspended at the above density. Control (unstained) cells were stained with just the fluorescent viability dye. Flow cytometry was performed to analyze the stained and control cells on a MACSQuant Analyzer, and the acquired data were further analyzed and plotted using FlowJo. At least 30,000 cells were acquired from each sample.

### Quantitative phase imaging and analysis

We acquired quantitative phase images of cells by suspending 10-15 μL of cell suspension from cultures at a density of 1×10^6^ cells/mL sandwiched between coverslips separated by a secure seal spacer (Invitrogen). We employed diffraction phase microscopy of our 3M system described previously for acquiring interferograms [33]. The raw interferograms were processed using inverse Fourier transform method to reveal phase images. We performed cell segmentation for confining only one cell in a frame for quantitative analysis and obtained 2294 images from the cells in all the five groups. We performed QPI measurements from three replicates of the same culture and also from the three different cultures of each cell type. The B cells were measured from three different donors. To understand the differences in the morphological differences between the cell types, we employed the recently developed VAMPIRE tool to analyze the QPI images [35, 36].

### Raman spectroscopy and analysis

We employed the 3M system, described previously, with 785 nm excitation for the acquisition of Raman spectra [33]. The cell samples were centrifuged and resuspended in media in PBS before transferring them to quartz cover glass for Raman measurements. For the cell lines, around twenty measurements were performed from each cell culture with three replicas to account for the variation in different measurements from the same culture. Moreover, cells were also measured from three different cultures. Similarly, the spectra from healthy B cells were recorded from cells from three different donors. We collected Raman spectra from cells in the Control (n=123), REH (n=198), RS4;11 (n=200), MN60 (n=202), and BALL-1 (n=236) classes. The integration time for each Raman spectrum (two acquisitions at 30s exposure) was 60 seconds. All the Raman data analyses were performed in MATLAB environment (Mathworks) unless otherwise noted. The spectra in the fingerprint region (700–1750 cm^−1^) were subjected to background removal using an iterative fifth order polynomial subtraction method [41]. The background corrected spectra were subjected to peak normalization using the maximum intensity of each spectra. We used principal component analysis for dimensionality reduction of the spectral dataset and identify a set of orthogonal abstract component spectra ordered according to the variance in the dataset they explain [42]. We used Orange data mining software to plot the scores of the principal components to visualize the clustering of the spectral data in the principal component space according to their cell type and their stage in leukemia progression [38]. We used TreeBagger class in MATLAB to implement Breiman’s original random forest algorithm for supervised classification of spectral dataset [43]. We included 100 trees in each random forest and performed 200 training iterations by selecting random subsets of spectral datasets in each iteration to ensure equal size of all the classes in the trained model and avoid bias due to unequal class sizes. The out-of-bag classification error rates were calculated for each trained model by subjecting the spectra that are intrinsically left out as part of random selection of examples during training the classifier.

## Acknowledgments

This work was partially supported by Connecticut Children’s, National Cancer Institute (R01 CA238025), the National Institute of Biomedical Imaging and Bioengineering (2-P41-EB015871-31), and the National Institute of General Medical Sciences (DP2GM128198). S.K.P. acknowledges the support of the SLAS Graduate Education Fellowship Grant. We also acknowledge the support from the Carole and Ray Neag Comprehensive Cancer Center, University of Connecticut School of Medicine. The schematic in Figure 1 was partially created with BioRender.com.

## References

[1] Woo JS, Alberti MO & Tirado CA (2014) Childhood B-acute lymphoblastic leukemia: A genetic update. Experimental Hematology & Oncology 3(1): 1–14.

[2] Good Z, et al (2018) Single-cell developmental classification of B cell precursor acute lymphoblastic leukemia at diagnosis reveals predictors of relapse. Nat Med 24(4): 474.

[3] Nordlund J, et al (2015) DNA methylation-based subtype prediction for pediatric acute lymphoblastic leukemia. Clinical Epigenetics 7(1): 11.

[4] Ross ME, et al (2003) Classification of pediatric acute lymphoblastic leukemia by gene expression profiling. Blood 102(8): 2951–2959.

[5] Lam VK, et al (2019) Machine learning with optical phase signatures for phenotypic profiling of cell lines. Cytometry 95(7): 757–768.

[6] Winnard Jr PT, et al (2017) Organ-specific isogenic metastatic breast cancer cell lines exhibit distinct Raman spectral signatures and metabolomes. Oncotarget 8(12): 20266.

[7] Basu S, Kolouri S & Rohde GK (2014) Detecting and visualizing cell phenotype differences from microscopy images using transport-based morphometry. Proceedings of the National Academy of Sciences 111(9): 3448–3453.

[8] Di Z, et al (2014) Ultra high content image analysis and phenotype profiling of 3D cultured micro-tissues. PloS One 9(10): e109688.

[9] Chhetri RK, Phillips ZF, Troester MA & Oldenburg AL (2012) Longitudinal study of mammary epithelial and fibroblast co-cultures using optical coherence tomography reveals morphological hallmarks of pre-malignancy. PLoS One 7(11): e49148.

[10] Boustany NN, Boppart SA & Backman V (2010) Microscopic imaging and spectroscopy with scattered light. Annu Rev Biomed Eng 12: 285–314.

[11] Ahmad A, et al (2018) Quantitative phase microscopy of red blood cells during planar trapping and propulsion. Lab on a Chip 18(19): 3025–3036.

[12] Karandikar SH, et al (2019) Reagent-free and rapid assessment of T cell activation state using diffraction phase microscopy and deep learning. Anal Chem 91(5): 3405–3411.

[13] Kasprowicz R, Suman R & O’Toole P (2017) Characterising live cell behaviour: Traditional label-free and quantitative phase imaging approaches. Int J Biochem Cell Biol 84: 89–95.

[14] Pavillon N, Hobro AJ, Akira S & Smith NI (2018) Noninvasive detection of macrophage activation with single-cell resolution through machine learning. Proceedings of the National Academy of Sciences 115(12): E2676–E2685.

[15] Park Y, Depeursinge C & Popescu G (2018) Quantitative phase imaging in biomedicine. Nature Photonics 12(10): 578–589.

[16] Jo Y, et al (2018) Quantitative phase imaging and artificial intelligence: A review. IEEE Journal of Selected Topics in Quantum Electronics 25(1): 1–14.

[17] Kim K, Kim KS, Park H, Ye JC & Park Y (2013) Real-time visualization of 3-D dynamic microscopic objects using optical diffraction tomography. Optics Express 21(26): 32269–32278.

[18] Kim K, et al (2017) Correlative three-dimensional fluorescence and refractive index tomography: Bridging the gap between molecular specificity and quantitative bioimaging. Biomedical Optics Express 8(12): 5688–5697.

[19] Kim K, et al (2013) High-resolution three-dimensional imaging of red blood cells parasitized by plasmodium falciparum and in situ hemozoin crystals using optical diffraction tomography. J Biomed Opt 19(1): 011005.

[20] Popescu G, et al (2008) Optical imaging of cell mass and growth dynamics. American Journal of Physiology-Cell Physiology 295(2): C538–C544.

[21] Lee S, et al (2017) Refractive index tomograms and dynamic membrane fluctuations of red blood cells from patients with diabetes mellitus. Scientific Reports 7(1): 1–11.

[22] Hosseini P, et al (2016) Cellular normoxic biophysical markers of hydroxyurea treatment in sickle cell disease. Proceedings of the National Academy of Sciences 113(34): 9527–9532.

[23] Paidi SK, Pandey R & Barman I (2020) Chapter 18 - emerging trends in biomedical imaging and disease diagnosis using Raman spectroscopy. Molecular and Laser Spectroscopy: 623–652.

[24] Paidi SK, Pandey R & Barman I (2020) Medical applications of Raman spectroscopy. Encyclopedia of Analytical Chemistry: 1–21.

[25] Bergholt MS, Serio A & Albro MB (2019) Raman imaging: Guiding light for the extracellular matrix. Frontiers in Bioengineering and Biotechnology 7: 303.

[26] Ember KJ, et al (2017) Raman spectroscopy and regenerative medicine: A review. NPJ Regenerative Medicine 2(1): 12.

[27] Schie IW, et al (2018) High-throughput screening Raman spectroscopy platform for label-free cellomics. Anal Chem 90(3): 2023–2030.

[28] Managò S, et al (2016) A reliable Raman-spectroscopy-based approach for diagnosis, classification and follow-up of B-cell acute lymphoblastic leukemia. Scientific Reports 6: 24821.

[29] Kumamoto Y, et al (2019) High-throughput cell imaging and classification by narrowband and low-spectral-resolution Raman microscopy. The Journal of Physical Chemistry B 123(12): 2654–2661.

[30] El-Mashtoly SF, et al (2015) Label-free Raman spectroscopic imaging monitors the integral physiologically relevant drug responses in cancer cells. Anal Chem 87(14): 7297–7304.

[31] Okada M, et al (2012) Label-free Raman observation of cytochrome c dynamics during apoptosis. Proceedings of the National Academy of Sciences 109(1): 28–32.

[32] Hamada K, et al (2008) Raman microscopy for dynamic molecular imaging of living cells. J Biomed Opt 13(4): 044027.

[33] Pandey R, et al (2019) Integration of diffraction phase microscopy and Raman imaging for label-free morpho-molecular assessment of live cells. Journal of Biophotonics 12(4): e201800291.

[34] Paidi SK, et al (2021) Coarse Raman and optical diffraction tomographic imaging enable label-free phenotyping of isogenic breast cancer cells of varying metastatic potential. Biosensors and Bioelectronics: 112863.

[35] Wu P, et al (2015) Evolution of cellular morpho-phenotypes in cancer metastasis. Scientific Reports 5(1): 1–10.

[36] Phillip JM, Han K, Chen W, Wirtz D & Wu P (2021) A robust unsupervised machine-learning method to quantify the morphological heterogeneity of cells and nuclei. Nature Protocols

[37] Movasaghi Z, Rehman S & Rehman IU (2007) Raman spectroscopy of biological tissues. Applied Spectroscopy Reviews 42(5): 493–541.

[38] Demšar J, et al (2013) Orange: Data mining toolbox in python. The Journal of Machine Learning Research 14(1): 2349–2353.

[39] Shu X, et al (2020) Artificial intelligence enabled reagent-free imaging hematology analyzer. arXiv Preprint arXiv:2012.08518

[40] Ayyappan V, et al (2020) Identification and staging of B-cell acute lymphoblastic leukemia using quantitative phase imaging and machine learning. ACS Sensors 5(10): 3281–3289.

[41] Lieber CA & Mahadevan-Jansen A (2011) Automated method for subtraction of fluorescence from biological Raman spectra. Appl Spectrosc 57

[42] Jolliffe I (2002) Principal component analysis, (Wiley Online Library,

[43] Breiman L (2001) Random forests. Mach Learning 45(1): 5–32.

