## Supporting Information for "Reagent-free Raman and quantitative phase imaging offer a unique morpho-molecular platform for recognition of malignancy and stages of B-cell acute lymphoblastic leukemia"

The authors disclose no potential conflicts of interest.

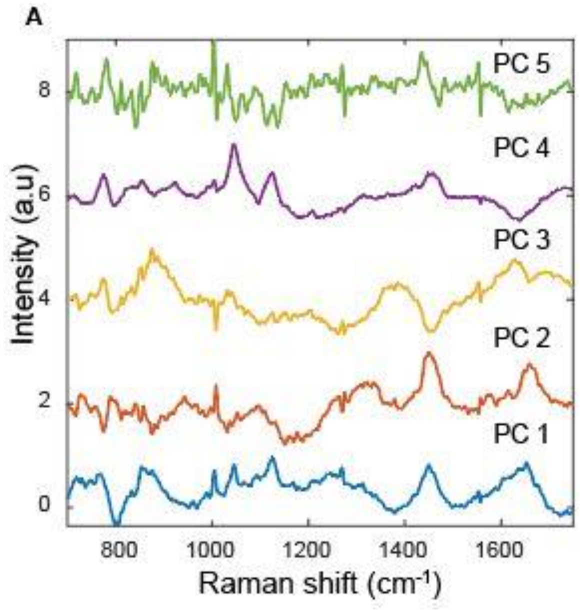


**Figure S1.** The first five principal component loadings obtained using principal component analysis of Raman spectra in the current study are shown.
